# Modeling membrane geometries implicitly in Rosetta

**DOI:** 10.1101/2023.08.23.554394

**Authors:** Hope M. Woods, Julia Koehler Leman, Jens Meiler

**Author notes:** Corresponding author: Jens Meiler, PhD, Department of Chemistry, Vanderbilt University, 21^st^ Ave S, Nashville, TN 37235 and Institute for Drug Discovery, Leipzig University Medical Faculty, Liebigstr. 19, 04103 Leipzig.

## Abstract

Interactions between membrane proteins (MPs) and lipid bilayers are critical for many cellular functions. In the Rosetta molecular modeling suite, the implicit membrane energy function is based on a “slab” model, which represent the membrane as a flat bilayer. However, in nature membranes often have a curvature that is important for function and/or stability. Even more prevalent, in structural biology research MPs are reconstituted in model membrane systems such as micelles, bicelles, nanodiscs, or liposomes. Thus, we have modified the existing membrane energy potentials within the RosettaMP framework to allow users to model MPs in different membrane geometries. We show that these modifications can be utilized in core applications within Rosetta such as structure refinement, protein-protein docking, and protein design. For MPs structures found in curved membranes, refining these structures in curved, implicit membranes produces higher quality models with structures closer to experimentally determined structures. For MP systems embedded in multiple membranes, representing both membranes results in more favorable scores compared to only representing one of the membranes. Modeling MPs in geometries mimicking the membrane model system used in structure determination can improve model quality and model discrimination.

## Introduction

Membrane proteins (MPs) exist in complex and diverse membrane environments in which they function. Cellular membranes adopt different geometries and cover a variety of different lipid compositions which affect membrane thickness. Lipid composition, thickness and curvature vary depending on species, cell type, and organelle the membrane belongs to [1–3]. Membranes also have local variability in lipid composition and thickness within the same bilayer. For example, lipid rafts can have a different lipid composition or thickness than their surrounding membrane. Protrusions and invaginations of the membrane, such as filopodia or caveolae, result from local membrane curvature in a larger bilayer. Vesicles, important for transport of various cargo molecules, are another classic example of curved membranes.

Interactions between MPs and their lipid bilayer affect one another’s shape, stability, and function [3–5]. Some MPs modify the membrane they are in, either by changing their thickness (for example some GPCRs [6]), curvature (for example BAR domains [7]), lipid composition (for example flippases, floppases, and scramblases [8]) or recruiting specific lipids to their location (for example some channels such as aquaporin Z (AqpZ) and the ammonia channel (AmtB) [9]). Reversely, the membrane bilayer can directly affect the function of many MPs such as in ABC transporters, RTKs, and mechanosensitive channels including some channels from the Piezo, TRP, MscL, and TREK families [2, 10–16].

Certain MPs such as piezo channels and BAR domains can induce membrane curvature [17, 18]. Curved membrane environments can impact stability and structure of MPs [19–23]. For example, bacteriorhodopsin’s stability depends on the degree of curvature of the membrane system [20]. MPs can also introduce pores into the membrane bilayers either by destabilizing the bilayer [24] or through containing a pore in their structure, as is the case for channels and transporters. Further, some MPs are large complexes that insert into two membranes in close proximity, such as gap junction channels across two cells [25] or efflux pumps that are multi-protein complexes that traverse the periplasm and insert into both the outer and inner membrane in Gram-negative bacteria [26].

Because structure determination of MPs in native membranes is practically challenging, MPs are typically solubilized into model membrane systems that allow for structure determination. These systems involve artificial membrane geometries that are rarely or never seen in cellular environments. Commonly used model membrane systems include detergent micelles, mixed detergent and lipid bicelles, lipid nanodiscs, and liposomes which vary in molecular composition [27, 28]. The choice of model membrane systems can impact the structure and stability of the MP [27, 29, 30]. Unfortunately, the model membrane system geometry in which experiments are performed is often disregarded during structure determination and is instead replaced by a flat bilayer (see below).

Despite significant progress in recent years, MP structure prediction and design lags behind that of soluble proteins [31]. Recent machine learning algorithms for protein structure prediction and design, such as AlphaFold2 [32, 33] and ProteinMPNN [34], neglect the membrane environment, which may cause inaccurate predictions in some cases [35]. Because state-of-the-art machine learning techniques depend on large amounts of training data, the vast under-representation of MP structures in the Protein Data Bank (PDB) as compared to soluble proteins [36, 37] (2.7% vs. 97.8% or 36-fold) is most likely the main reason for the limitations of these methods when it comes to MPs.

Computational modeling of membrane bilayers is achieved using explicit or implicit solvent. Molecular dynamics simulations often use explicit solvent models where diverse lipid bilayers or model membrane systems are modeled by representing individual lipid or detergent molecules, which is computationally expensive [38]. In contrast, implicit membrane energy functions, like those used in Monte-Carlo techniques such as the Rosetta software, are computationally efficient and model the effects the membrane has on protein structures [31, 39, 40] through the interaction of the protein with a continuous medium of average bilayer properties. Implicit membrane models such as IMM1 [40] are useful for many applications, including MP structure prediction, protein-protein docking, and design [39, 41–43]. While much progress has been made in improving these models to more accurately represent the membrane environment, most implicit membrane models are limited to a flat representation of the membrane, or a “slab” model. The slab neglects the fact that membrane proteins may exist and function in different geometries such as highly curved membranes with a radius of curvature as low as 50 Å [12], model membrane systems such as micelles, bicelles, nanodiscs, or double membranes.

Recent improvements to the Rosetta implicit membrane model allow more realistic bilayer representations by customizing parameters to model different lipid compositions [39]. This update also included the ability to model aqueous pores in MPs. One limitation of this membrane model is the lack of geometrical diversity it is able to simulate. One method and web server, PPM 3.0 ("Positioning of Proteins in Membranes"), accounts for different membrane geometries such as curvature or two membranes [44]. The PPM 3.0 server evaluates whether a protein is more likely to exist in a flat or curved membrane based on the transfer free energy from water to the membrane environment, calculated from the protein structure. The PPM server runs a grid search of a MP placed in a flat membrane, a curved membrane, and also testing different radii of curvature from 80 to 600 Å, to find the minimum transfer free energy and therefore the ideal membrane geometry for the protein structure in question. In case the MP is embedded in two or more membranes, it can give the optimal relative placement of each bilayer around the protein. The authors provide many examples of MP structures that were optimally placed in curved membranes or in multiple membranes [44]. This algorithm was incorporated into the PPM webserver and OPM (“Orientations of Proteins in Membranes”) database. Thus one can input their protein structure of interest or look up the PDB ID to see the predicted optimal membrane placement [45, 46]. FMAP ("Folding of Membrane Associated Peptides"), uses the peptide sequence to predict helical propensity, membrane depth, and ideal membrane geometry for different environments (water, lipid bilayer, or a micelle with a fixed radius depending on detergent [47]. Both methods are limited to placing the protein into the membrane and do not allow for further modeling.

Here we introduce a substantial code refactor within the RosettaMP framework to allow the use of different geometries in the implicit membrane model and in modeling applications. These geometries include the traditional flat model, an ellipsoid model to represent model membrane system geometries such as micelles, bicelles, and nanodiscs, a spherical bilayer similar to a vesicle to simulate curvature, and a model with two membranes. We provide examples of how using these models improve computational predictions for protein design, protein-protein docking, and high-resolution refinement.

## Results

### We added different implicit membrane geometries to RosettaMP

This substantial refactor was geared to achieve multiple goals such as: (1) adapting the current implicit membrane models for geometries that mimic micelles, bicelles, vesicles, membrane curvature, and double bilayers, (2) adapting the code interface to facilitate implementation of new geometries (for example lipidic cubic phases), (3) streamlining the current implementation by disentangling the pore from membrane geometries to be used independently and creating a membrane representation from a single, consistent implementation, thereby (4) minimizing code duplication, (5) improving testing, and (6) visualizing membrane geometries onto protein structures.

### Previous implicit membranes in Rosetta only describe a flat membrane

Mathematically, the membrane is modeled by a transition function that describes the transition from the aqueous environment to the hydrophobic environment of the membrane. Two transition functions had previously been implemented, one based on IMM1 and an adaptation to model different lipid compositions. Disregarding the pore, both transition functions score atoms based on their distance from the center of a membrane plane, meaning that the hydrophobicity at a specific position is only dependent on the depth in the membrane (z-coordinate). This creates the implicit slab membrane as shown in Figure 1A.

**Figure 1:**
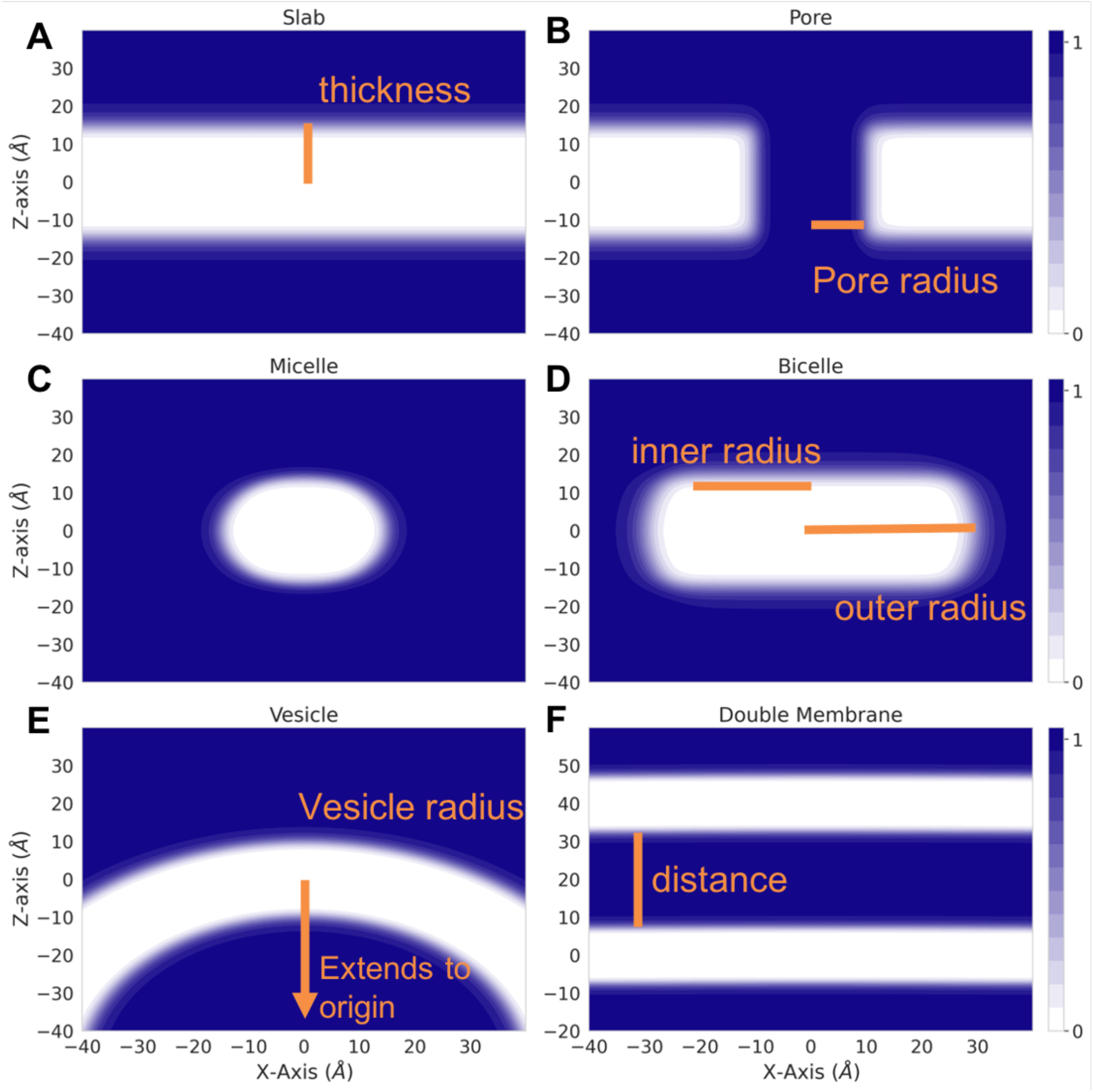
Implicit Membrane Energy Transition Functions. Value of transition functions for different geometries are shown in the XZ plane, at Y = 0. Atoms with transition function values of 0 (white) are evaluated as being in a completely hydrophobic environment. Atoms with transition functions values of 1 (blue) are evaluated as being in a hydrophilic environment. Atoms with transition function values in between are weighted based on their depth in the mebrane. Input parameters are shown in orange. A. Slab B. Slab including a pore. C. Micelle, inner radius = 0 D. Bicelle E. Vesicle F. Double Vesicle

### Expanding beyond flat membranes through implementation of mathematical representations of desired geometries

To model different geometries, the new transition functions must be dependent on the distance to the membrane center in more than just one dimension. The pore was the first adaptation that introduced a three-dimensional dependence (Figure 1B). In order to create an ellipsoid geometry similar to micelles and bicelles, we created a functional form that combines the slab transition function [48] with a function that describes the dependence on the distance to the center of the ellipsoid (Figure 1C and D). Although micelles and bicelles have different chemical and physical properties, we use the same function to describe their shape whereas the micelle is a special case of the 7icelles with an inner radius of the ellipsoid (Figure 1D) just large enough to surround the transmembrane region of the protein. To model a curved membrane such as a vesicle, we model the membrane as a sphere around an origin (0, 0, -z) and keep the membrane center as the location of the membrane residue [41]. The vesicle concept is extended to describe two membranes by a double vesicle (two concentric spheres) that, with a large radius, represent two essentially flat membranes (Figure 1F). Detailed descriptions of equations and parameters can be found in the Methods section.

### Code design ensures current and new geometries are compatible across membrane energy functions

To achieve our goal of generalizability across score functions and simplify the implementation of new geometries, we introduced an abstract base class into the RosettaMP framework that all other membrane geometry classes are derived from (Figure S1). By defining pure virtual functions within the MembraneGeometry base class, we ensure that all derived classes (or new geometries) define the transition function and the derivative of the transition function for that geometry. Functions used across different geometry classes are defined in the MembraneGeometry base class to decrease code duplication. Implementation details, function definitions, and score terms that depend on them, can be found in the Methods section. This implementation ensures that new geometries can be implemented by solely creating a class and updating options for setting the new geometry without updating each individual score term that depends on the transition function. This also allows for geometries to be used across score functions, and the decoupling of the pore model from the membrane score terms. Therefore, membrane dependent score terms implemented before the pore can utilize the pore model as well.

### Optimized code design allows for integrating membrane geometries into different applications

Implementation of the membrane geometries following the same object oriented design principles used in the RosettaMP framework allows for use in any application accessible to the RosettaMP framework [41], including but not limited to design, refinement, and protein-protein docking. This implementation also ensures a consistent user interface with the same command line options for setting geometry parameters across applications. The membrane geometries are also accessible through the different Rosetta scripting interfaces, including RosettaScripts [49] and PyRosetta [50].

### New implementation decouples pore model from energy functions, as tested for design

Alford et al. described an improvement in protein design using their newly developed energy function, *franklin2019*, compared to previous membrane energy functions [39]. However, it was not clear if these improvements came from the new energy function, from the inclusion of the pore, or a combination of the two. This could not be tested because the pore was incompatible with older membrane energy functions. Our reimplementation of the pore function into the MembraneGeometry class allows for its use across applications and energy functions. Therefore, we are able to test protein design including the pore for both energy functions, *mpframework2012* and *franklin2019*. We used an established membrane protein design benchmark set to test where this improvement comes from [51]. The test uses three different metrics to describe performance: (1) sequence recovery is the fraction of native residues recovered after design over all designable positions. (2) non-random recovery of individual amino acids, where the recovery rate for each amino acid is calculated relative to the background probability of randomly guessing the native amino acid (1/20). (3) The Kullback-Leibler (KL) divergence measures how different distributions of designed amino acids are compared to native distributions. The design test done on the subset of proteins from the dataset that contain a pore to highlight the effect of accounting for an aqueous pore. The design protocol was run with four different energy function conditions: *mpframework2012* without the pore included, *mpframework2012* with the pore, *franklin2019* without the pore, and *franklin2019* with the pore. When analyzing all residues in each protein, *franklin2019* outperforms the *mpframework2012* energy function with respect to all three metrics. Inclusion of the pore in either *franklin2019* or *mpframework2012* score functions does not improve the sequence recovery or non-random recovery (Figure 2A, B). If only pore-facing residues are taken into account, inclusion of the pore for *franklin2019* shows a significant difference in all three metrics. Differences in performance when including the pore for *mpframwork2012* are minor. An improvement in KL divergence is seen with *franklin2019* across all residues, and the difference is greater when only considering pore-facing residues (Figure 2C). On the other hand, KL divergence does not show an improvement for *mpframework2012* when including the pore.

**Figure 2:**
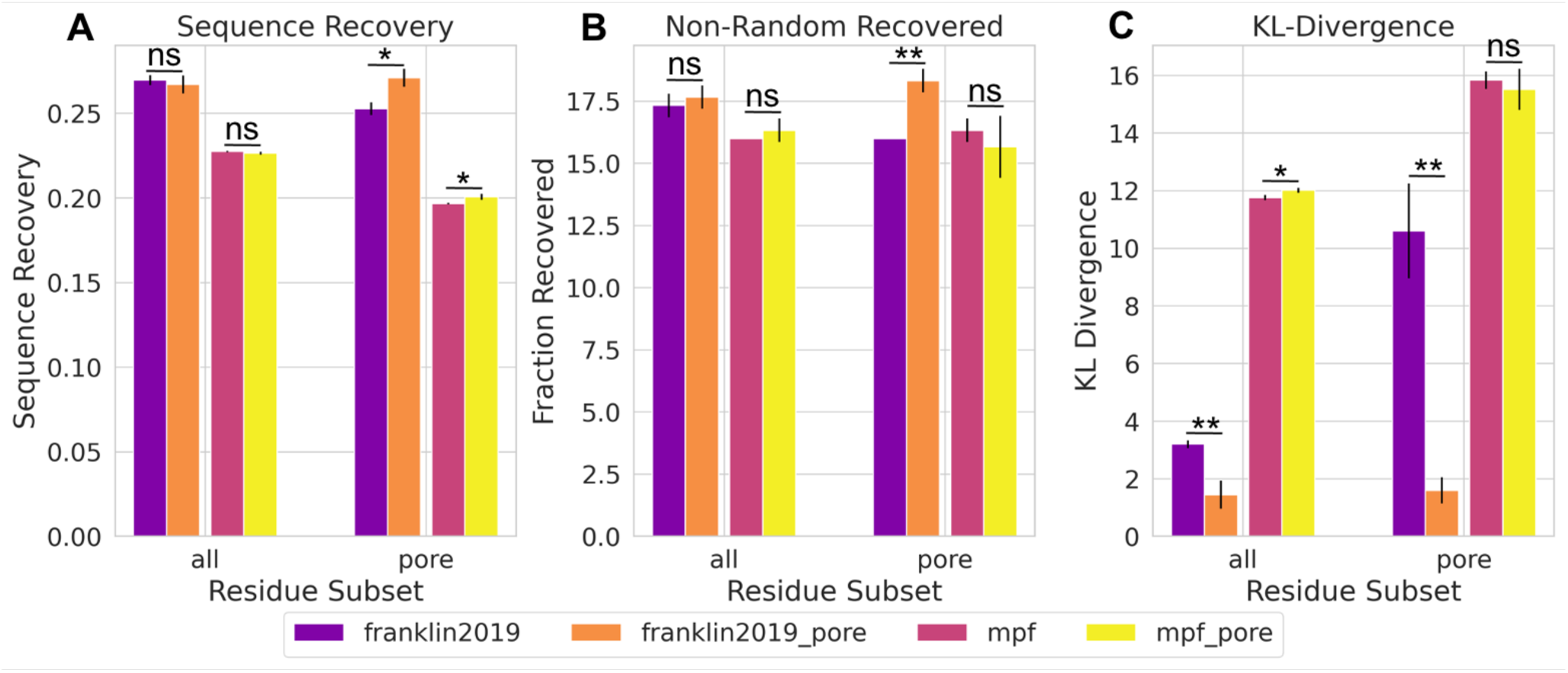
Sequence Design of Pore Across Score Functions. Our implementation allows the decoupling of the pore model from the energy functions, here shown for franklin2019 and mpframework2012 (mpf). This data shows that the improvements for design originate in the franklin2019 energy function where the pore model only plays a minor role in the improvements. A) Fraction of native residues recovered after design for all residues (left) and pore-facing residues (right). Higher values are expected for more accurate energy functions. Results using the franklin2019 function are purple, franklin2019 with the pore are orange, mpframework2012 are pink, mpframework2012 are yellow. B) Average recovery rates for each individual amino acids relative to the background probability of randomly guessing the native amino acid (1/20). Higher values are expected for more accurate energy functions. C) Kullback-Leibler (KL) divergence measures how different distributions are of designed amino acids compared to native distributions. Lower values are expected for more accurate energy functions.

### Refinement of structures in a curved membrane results in higher quality models

Several MPs exist in curved membranes. To take curvature into account we have implemented a vesicle geometry where the user would set the vesicle radius to define the degree of curvature. The vesicle radius is the distance from the center of the vesicle to the center of the membrane. We tested refinement in a curved membrane on the mechanosensitive channel Piezo 1 (PDB ID 6B3R). The curve of the detergent micelle around Piezo 1 can clearly be seen in the cryo-EM images (Figure 2 in [17]). When the slab transition function is mapped onto the structure it is clear that it does not properly represent the hydrophobic environment needed (Figure 3A). The result of running high-resolution refinement with this poor membrane placement is evident from the large structural changes in the output models as compared to the input structure (Figure 3B, C, D), refinement essentially breaks up the protein. Even the lowest scoring model with the slab geometry has an RMSD of 26 angstroms with large changes compared to the starting structure shown in gray (Figure 3B, C, D). The vesicle transition function with a radius of 120 Å (based on PPM 3.0 predictions) better fits the membrane geometry in Piezo 1 (Figure 3E). The lowest scoring model is much closer to the starting structure for the vesicle geometry (∼8A, Figure 3F and G). While both geometries also produce models that have large positive scores due to clashes, the percentage of low-scoring, high-quality models is much higher for the vesicle geometry than for the slab (Figure 3D and H). In another curved membrane example of potassium chloride cotransporter KCC2, the slab geometry also results in more high scoring, low-quality models (Figure S2 C and F). In this case though, the slab geometry still also produces several low RMSD, low scoring models, potentially because the difference in the implicit membrane from the slab and vesicle geometries are not as great as for Piezo 1 (Figure S2 A and D).

**Figure 3:**
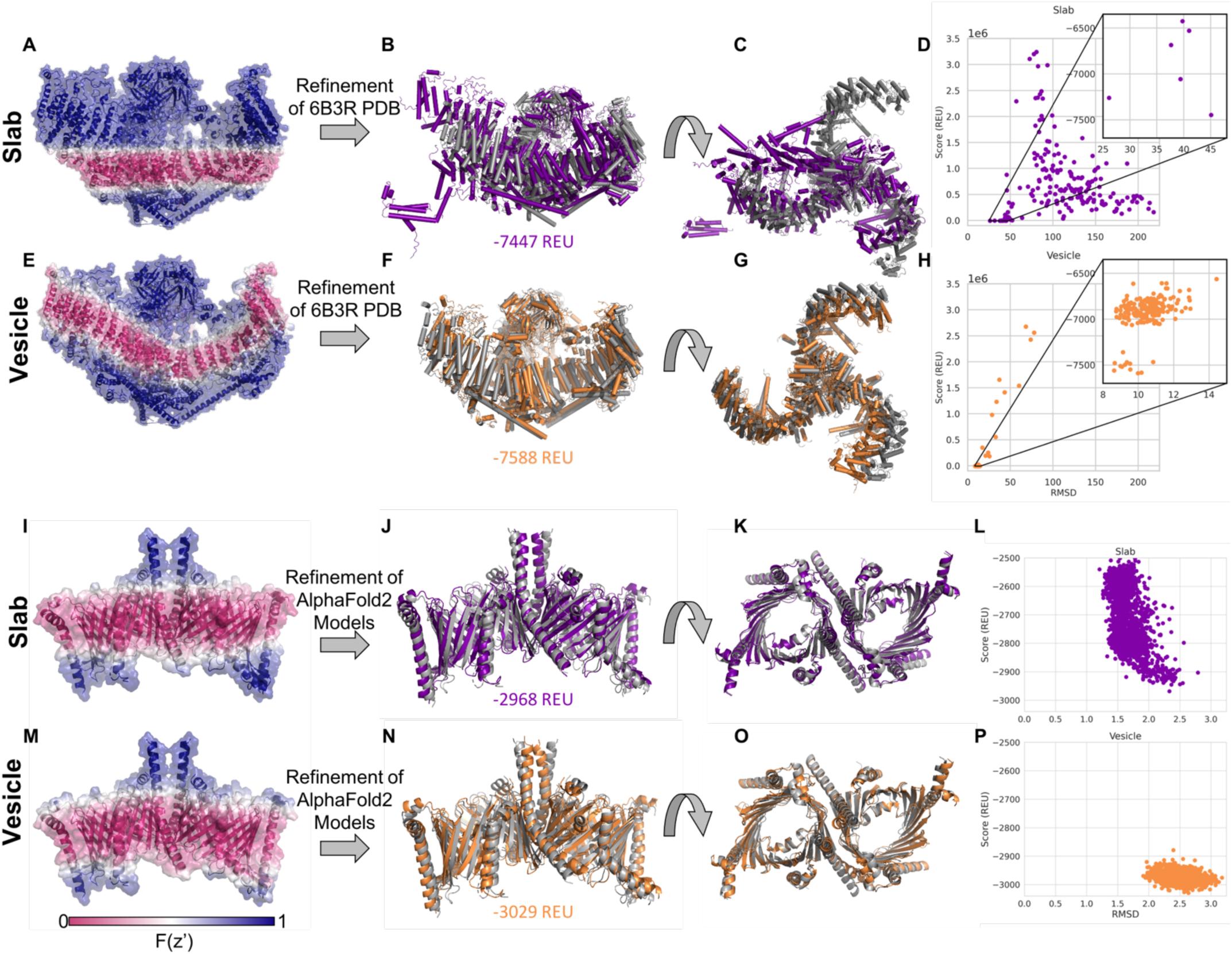
Refinement into vesicle geometries. A) Mechanosensitive channel Piezo 1 (PDB ID 6B3R) with the slab transition function mapped onto the structure. A value of zero is shown in pink and indicates that residue is scored in a hydrophobic environment. A value of one is show in blue and indicates that residue is scored in an aqueous environment. B) Lowest scoring structure after refinement with slab geometry ran on PDB structure with a score of -7447 Rosetta Energy Units (REU). Starting structure (PDB ID 6B3R) shown in gray C) Top view of structure in B. D) RMSD with respect to PDB 6B3R vs score of models from refinement with slab geometry, inset plot shows lowest scoring models. E) Same structure as A with Vesicle geometry transition function with a radius of 120 Å. F) Lowest scoring structure after refinement with vesicle geometry ran on PDB structure with a score of -7588 REU G) Top view of structure in F. H) RMSD with respect to PDB 6B3R vs score of models from refinement with vesicle geometry, inset plot shows lowest scoring models. I) Mitochondrial TOM complex (PDB 6UCU) with the slab transition function mapped onto the structure. J) Lowest scoring structure from generating structures with AlphaFold2 followed by refinement in Rosetta with slab geometry ran on PDB structure with a score of -2968 (REU). Starting structure (PDB ID 6UCU) shown in gray K) Top view of structure in J. L) RMSD with respect to PDB 6UCU vs score of models from AlphaFold2 followed by refinement with slab geometry M)) Same structure as I with Vesicle geometry transition function with a radius of 180Å. N)) Lowest scoring structure from generating structures with AlphaFold2 followed by refinement in Rosetta with slab geometry ran on PDB structure with a score of -3029 (REU). O) Top view of structure in N. P) RMSD with respect to PDB 6UCU vs score of models from AlphaFold2 followed by refinement with vesicle geometry

### Refinement and scoring of AlphaFold2 models in a curved membrane

The TOM complex (translocase of the outer membrane) was recently determined by cryo-EM (PDB ID 6UCU) [52]. PPM 3.0 predicts that TOM complexes induce membrane curvature with a radius of ∼180 Å [44]. How the TOM complex structure would be scored in the slab and vesicle geometries can be seen by mapping the transition function values onto the structure (Figure 3I, M). As a demonstration of how one could use this implementation in combination with other tools, we used AlphaFold2 multimer [53] to predict the structure of the TOM complex from the sequence. Then, we used the FastRelax protocol to optimize and score the predicted models in the slab and the vesicle implicit membrane geometries. The lowest scoring models for both slab (Figure 3J and K) and vesicle geometries (Figure 3N and O) are similar to the determined structure shown in gray. While the slab geometry results in output models with slightly lower RMSD values to the PDB structure, they have a much larger range of scores, indicating that the refinement can’t find an ideal fit of the flat membrane around the protein. Almost all the models from the vesicle geometry have lower scores than the models from the slab geometry (Figure 3L and P), indicating a better fit of the protein in the vesicle.

### Modeling two membranes provides a more accurate representation of membrane protein systems spanning different bilayers

Another limitation of previous implicit membrane energy functions is only being able to account for one membrane per system. While one membrane is often sufficient, this ignores cases where one protein complex spans two membranes, such as in gram negative bacteria. Here we introduce a double vesicle option that represents the membrane as two vesicles, sharing a single origin but with different radii. One can approximate two flat membranes by setting a large radius for the vesicles, such as 1000 A.

### Implicit membrane for lipid bilayers of two interacting cells

The first example we tested is a gap junction channel, where two transmembrane proteins in lipid bilayers of different cells are forming the gap junction. We used the connexin 46 gap junction channel (PDB ID 6MHQ) as a starting structure [25] and refined this structure in both the original slab membrane and in a double membrane. The value of the transition function was mapped onto the structure to visualize the membrane environment for each case (Figure 4A, B). The double membrane models all have lower scores than slab models (Figure 4E). This drop in score can be attributed to the *fa_water_to_bilayer* term which evaluates to zero when residues are sufficiently far from the membrane [39]. Therefore, including the second membrane results in more residues with negative *fa_water_to_bilayer* values. The output models are similar for both geometries and the lowest scoring models overlap well with the starting structure (Figure S3A).

**Figure 4:**
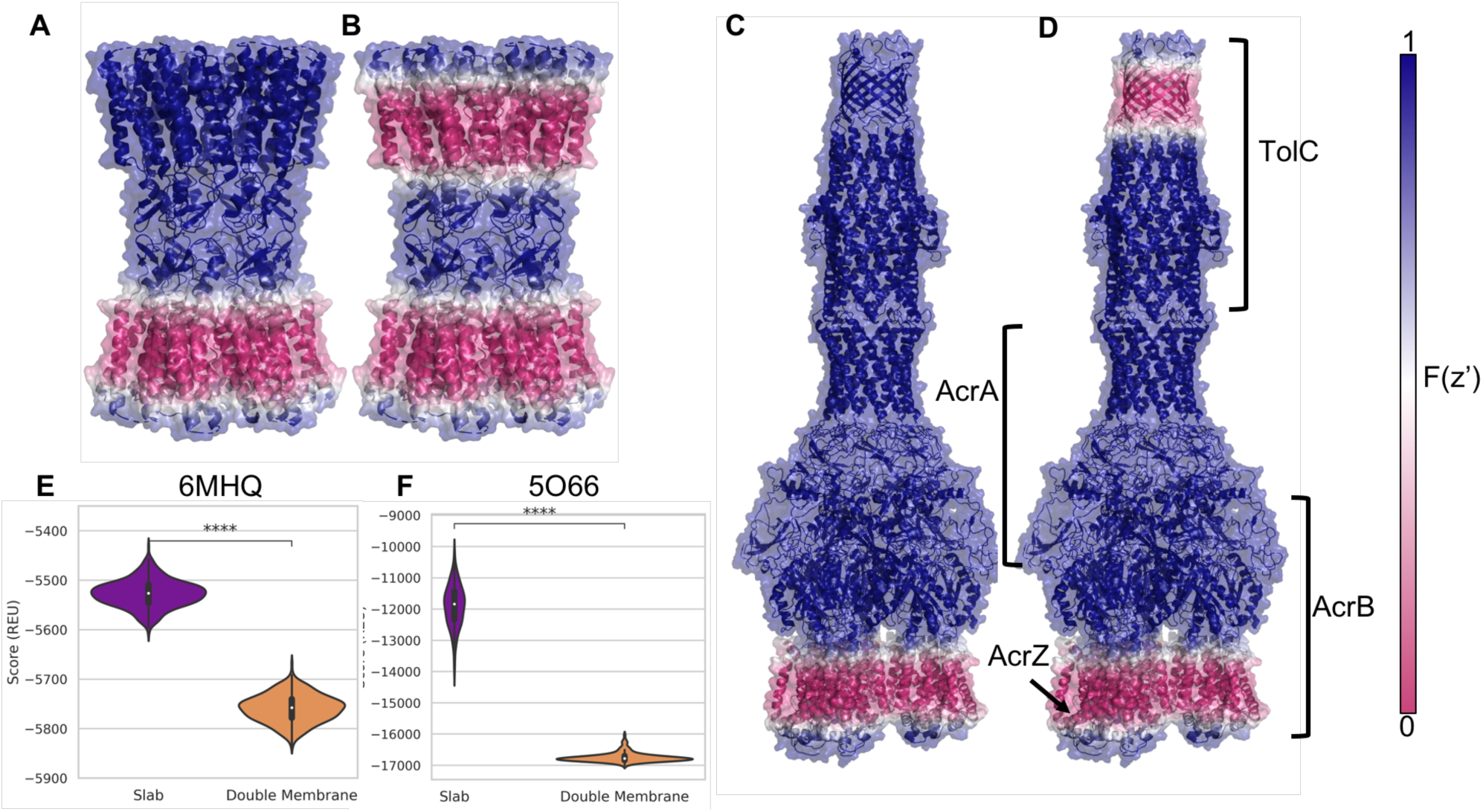
Refinement in double membrane. A) An intercellular gap junction channel (PDB ID 6MHQ) with the slab transition function mapped onto the structure. A value of zero is shown in pink and indicates that residue is scored in a hydrophobic environment. A value of one is show in blue and indicates that residue is scored in an aqueous environment. B) The same structure as in A with the double vesicle transition function mapped on to the structure. C) AcrABZ-TolC multidrug efflux pump (PDB ID 5O66) with the slab transition function mapped onto the structure. D) The same structure as in C with the double vesicle transition function mapped onto the structure. E) Score distributions for structural models created by running relax with the starting structure from PDB ID 6MHQ with franklin2019 in the slab (purple) and double vesicle (orange) geometries. F) Score distributions for structural models created by running relax with the starting structure from PDB 5O66.

### Implicit membranes for inner and outer membranes of gram-negative bacteria

The second example we tested was the AcrABZ-TolC efflux pump (PDB ID 5O66) [26]. The transition function values for the slab (Figure 4C) and the double vesicle (Figure 4D) are mapped onto the structure to visualize the membrane for each case. In this complex, TolC is embedded in the outer membrane, while the AcrB and AcrZ subunits are embedded in the inner membrane (Figure 4D). The AcrA subunit is in the periplasm [26]. Similar to the previous gap junction example, we see a drop in score with the double membrane simulations that can be partially attributed to more residues having a negative *fa_water_to_bilayer* scores since there is a second membrane (Figure 4F). Interestingly, the score distributions are much wider for models produced by slab geometry where there is a single membrane represented (Figure 4F and Figure S3C). This results from sampling larger changes of the membrane position relative to the structure in the slab geometry compared to the double vesicle geometry, which anchors the complex better in the membrane. The larger range of scores is caused by the larger range of values in the *fa_water_to_bilayer* score term for the slab models than in the double membrane models. The output models generated by both geometries superimpose well with the starting structure and a similar tilt of TolC with respect to the rest of the complex is observed for both geometries (Figure S3B).

### Incorporating experimental model membrane systems geometries in computation leads to more accurate models

For most structure determination methods, MPs must be extracted from the plasma membrane and reconstituted into a model membrane system. The composition and shape of the model membrane systems can impact the conformation of MPs. Having the ability to model MPs in geometries similar to the model membrane system in which the structure was determined in (or in which restraints were acquired) may provide additional insight into how model membrane systems introduce artifacts in MP structure and function. We use the ellipsoid model to capture the shape of model membrane systems including micelles and bicelles.

### Protein-protein docking in experimentally relevant membrane geometry leads to better model discrimination

To demonstrate how modeling MPs in model membrane system geometries can be useful, we ran the protein-protein docking protocol, MPdock [42], on the glycophorin A homodimer, each with a single membrane spanning domain [54]. The structure of glycophorin A was determined by NMR in detergent micelles [55]. The MPdock protocol consists of two parts: a pre-optimization step where the single chains are pulled apart and optimized, and the docking step. Each step was run using either the slab or micelle geometry. The transition function values for the slab and micelle geometries are mapped on the input structure (Figure 5A and D). The lowest scoring models for each geometry are superimposed onto the starting structure (PDB ID 1AFO) shown in gray (Figure 5B and E). The output models of the final step for each geometry are evaluated using the interface score versus the interface RMSD (Figure 5C and F), indicating that the models produced with the micelle geometry result in lower scores. The score function is also better able to discriminate lower RMSD models using the micelle geometry as indicated by the funnel metric (Pnear) value of 0.63 compared to 0.06 for the slab geometry. Pnear ranges from zero to one, where higher values indicate the metrics’ (in this case the interface score) ability to favor conformations with lower RMSD value [56].

**Figure 5:**
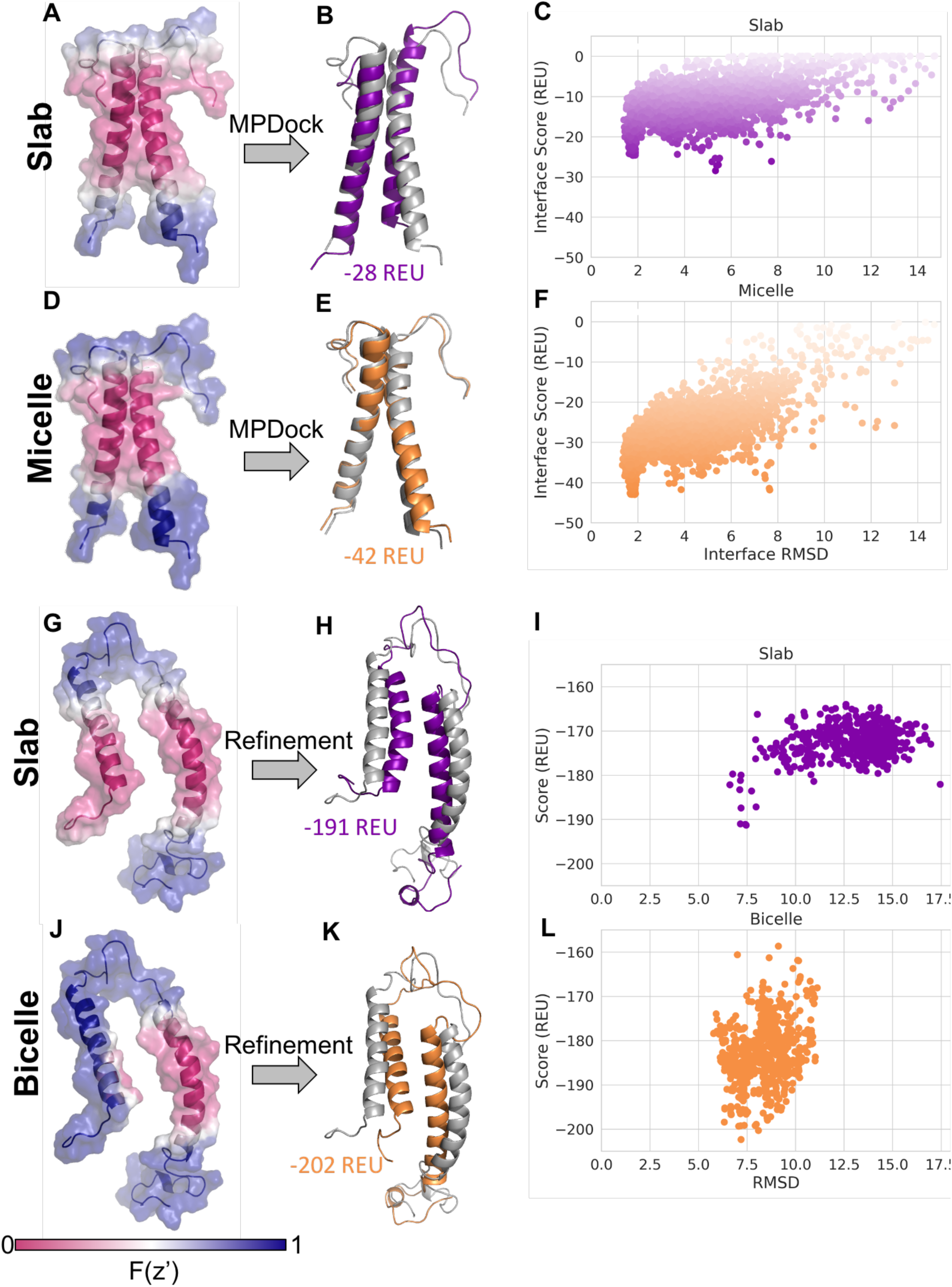
Docking and refinement with structures in model membrane systems. A) Glycophorin A dimer with slab transition function mapped onto the first NMR ensemble model from PDB ID 1AFO. B)) Lowest scoring structure after docking with slab geometry with an interface score of -28 REU. Starting structure, PDB 1AFO, shown in gray. C) Interface RMSD and interface score from running MPdock protocol for PDB ID 1AFO in slab geometry, pnear value of 0.06. D) The same structure in A with the micelle geometry with inner radius 1 Å transition function mapped onto the structure. E) Lowest scoring structure after docking with micelle geometry with an interface score of -42 REU. F) Interface RMSD and interface score from running MPdock protocol for PDB ID 1AFO in bicelle geometry, pnear value of 0.63. G) KCNE3 with slab transition function mapped onto the structure of the 4th conformation in NMR ensemble PDB ID 2NDJ. H) Lowest scoring structure after refinement with slab geometry with a score of -191 REU. Starting structure, PDB 2NDJ, shown in gray. I) RMSD with respect to starting structure vs score of models from refinement with slab geometry. J) The same structure in G with the bicelle geometry with inner radius 2 Å transition function mapped onto the structure K) Lowest scoring structure after refinement with bicelle geometry with a score of -202 REU. L) RMSD with respect to starting structure vs score of models from refinement with bicelle geometry.

### Refinement of transmembrane helix in membrane geometry similar to experiment results in models more similar to experimental structure

The structure of KCNE3 was determined with NMR in bicelles [57], showing a single span MP with an N-terminal amphipathic helix. However, when scoring the models in the NMR ensemble with the traditional slab membrane model, some conformations such as the fourth conformation in the ensemble, are scored as if the N-terminal amphipathic helix was dipping into the membrane (Figure 5G). We chose to work with the fourth conformation in the NMR ensemble to highlight how the ellipsoid geometry can be utilized to more accurately model the hydrophobic environment the protein is in. This scenario is more accurately modeled using the bicelle geometry, where the N-terminal helix is now scored as if it is resting on the edge of the bicelle (Figure 5J). In fact, when this conformation is refined and scored using the slab and bicelle geometries, the bicelle geometry results in lower scoring models and lower RMSD values relative to the starting structure (Figure 5I and L). The relative position of the two helices with respect to each other changes for both geometries, bringing the helices closer together, which can be seen with the lowest scoring structures (Figure 5H and 5K), resulting in RMSD values ∼7A.

### Representing geometry of model membrane systems allows separating artifacts from model membranes vs. structure determination method

While model membrane systems can impact the protein structure, even in similar systems different structure determination techniques can result in different conformations [58]. For example, the structure of outer membrane protein G (OmpG) has been determined in micelles with both NMR (DPC micelles) [59] and crystallography (OG micelles) [60]. We ran refinement on the NMR model and crystal structure in both slab and micelle geometries. The NMR model (PDB ID 2JQY) has a large loop that seemingly wraps around the micelle, but in the slab, geometry is treated like it is in the membrane (Figure 6A). In the micelle geometry with an inner radius of 8 Å this loop is scored like it is in an aqueous environment (Figure 6B). After refinement the models do not have large structural differences to the input structure (Figure 6C and D). However, the score distribution for the micelle geometry is shifted towards more negative scores as compared to the slab geometry (Figure 6D). Mapping the slab and micelle transition functions (micelle inner radius of 8 A) onto the crystal structure (PDB ID 2X9K) does not result in large visual differences (Figure 6E and F). Refinement in both geometries sample similar structures with scores in the same range (Figure 6G and H).

**Figure 6:**
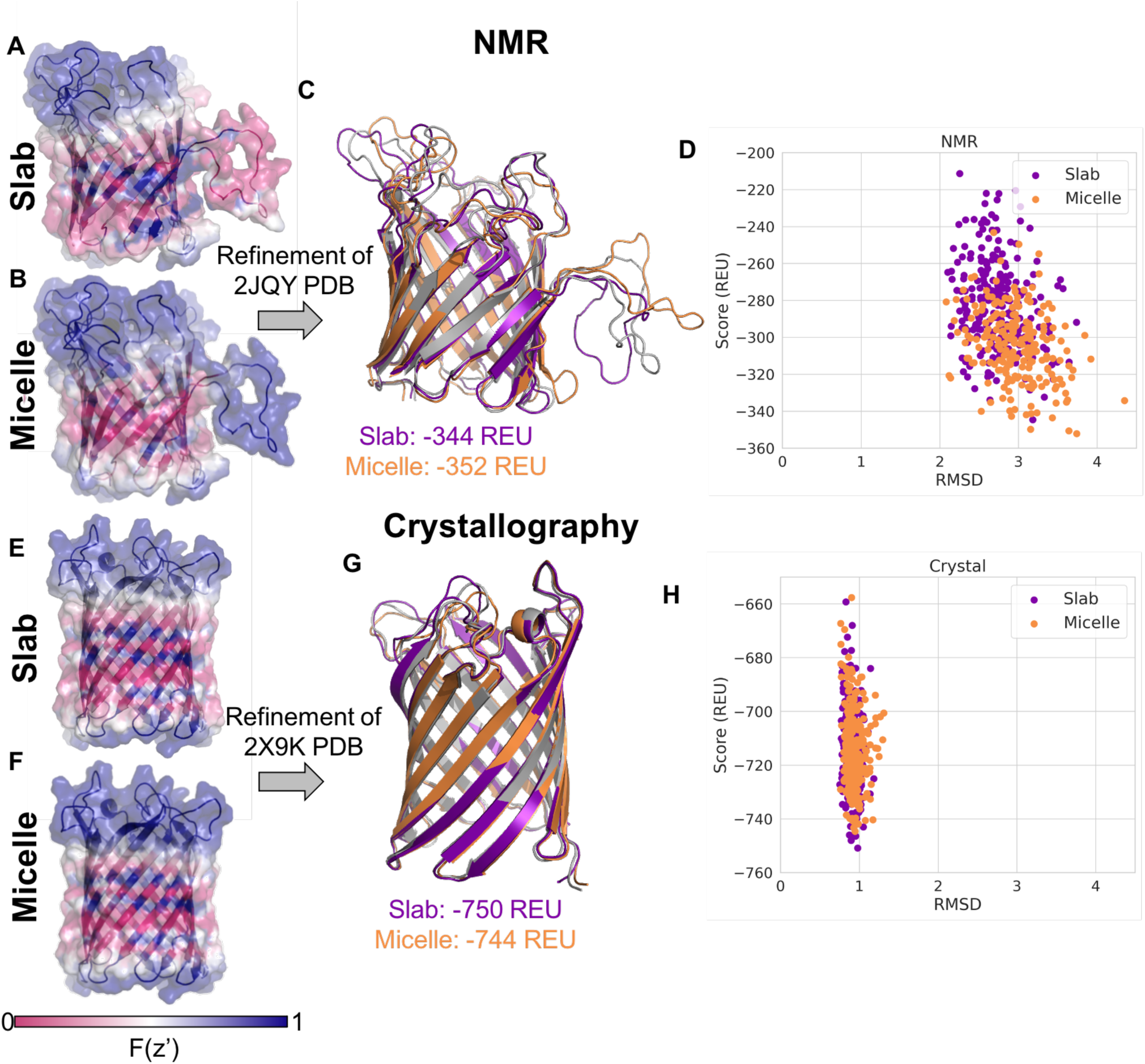
Refinement of OmpG NMR and Crystal Structures. A) OmpG with slab transition function mapped onto the structure of the 1st conformation from the NMR ensemble from PDB ID 2JQY. B) The same structure in A with the micelle geometry with inner radius 8 Å transition function mapped onto the structure. C) Lowest scoring structure after refinement, model from slab geometry is purple with a score of -344 REU, model from micelle geometry is orange with a score of -352 REU, starting structure (PDB ID 2JQY) in gray. D) RMSD with respect to starting structure vs score of models from refinement, slab geometry in purple, micelle geometry in orange. E) OmpG with slab transition function mapped onto the crystal structure PDB ID 2X9K. F) The same structure in E with the micelle geometry with inner radius 8 Å transition function mapped onto the structure. G) Lowest scoring structure after refinement, model from slab geometry is purple with a score of -750 REU, model from micelle geometry is orange with a score of -744 REU, starting structure (PDB ID 2X9K) in gray. H) RMSD with respect to starting structure vs score of models from refinement, slab geometry in purple, micelle geometry in orange.

## Discussion

We have implemented a general framework to adapt implicit membrane models to different geometries. Within this framework, there are currently four options available: slab, bicelle and micelle, vesicle, and two membranes. These geometries are compatible with existing score functions within the RosettaMP framework and allow users to accurately model MPs within specific environments based on their target.

The geometry framework was implemented to decouple the pore representation from the score functions, allowing older score functions within the MPFramework to be tested with the pore. We were able to analyze improvements of including the pore for both *franklin2019* and *mpframework2012* score functions. Sequence design metrics across all residues in the dataset only showed minimal differences when including the pore for either score function tested. However, inclusion of the pore only influences a handful of residues in each structure. When only considering pore-facing residues, there is a significant difference for all three metrics when structures are designed with *franklin2019*, yet the difference for the *mpframework2012* score function is minor. *Franklin2019* may perform better with the pore since it was parameterized and tested with the pore implemented. On the other hand, *franklin2019* may overestimate the favorability of residues being buried in the membrane [61]; therefore, scoring pore-facing residues as being in an aqueous environment would help compensate for that.

The vesicle geometry allows MPs to be modeled in a curved membrane, providing a more realistic representation of the MPs involved in membrane curvature. Membrane curvature may not be as important for small, compact MPs since the curvature would have to be high for the residues considered in the membrane to change from the slab to the vesicle representation. Large protein complexes are more likely to benefit from a curved membrane representation. When refining the Piezo 1 channel in both slab and vesicle, it is clear that the slab model produces low-quality output models where the protein essentially breaks apart. For the AlphaFold2 model of the TOM complex followed by Rosetta refinement, we see lower scoring and higher RMSD models for the simulations run with the curved membrane.

We can now model large complexes that span two membranes using a double vesicle geometry with a large radius. For these examples (gap junctions and ArcABZ-TolC complex), there is a clear drop in score for systems when they are scored with the additional membrane. This is expected since the membrane domain of these complexes would be unstable if not embedded in the membrane. For the examples tested, the refinement protocol produces relatively tight ensembles with small differences in the RMSD of models.

MP structures are often determined in a model membrane system. Although the conformations scientists may be interested in are the ones in the native membranes, we typically only have access to structures in model membrane systems. Model membrane systems are often a surrogate for native membranes during structure determination experiments, but it is important to keep in mind that the model membrane affects the energy landscape of the MP. Although the differences in scoring may be subtle, in some of the examples shown using the model membrane geometry resulted in lower scoring models. Being able to model different model membrane systems geometries might now allow us to separate effects from the model membrane system and the structure determination method, yet further studies are needed in this area.

In this work, we show how implicit membrane energy functions can be adopted to represent alternate membrane geometries instead of only a flat bilayer. Using appropriate geometries can improve both scoring and sampling. For most examples shown we observe lower scores when using the appropriate geometry. We observe narrower ranges in both RMSD and score of models being sampled when using the appropriate geometry, with the sampled conformations having lower RMSD, lower scores, or both indicating a more focused sampling of desired conformations.

## Conclusion

Modeling and designing MPs continues to be a challenge in the protein structure field. However, many advances are being made to improve computational predictions for MPs. The complex and diverse environments MPs are in, contributes to this challenge. The framework introduced here allows modeling MPs in different geometries that cover experimentally used model membrane systems, curved membranes, and two membranes. The ability to model MPs in different model membrane systems opens the door for studies to separate the effect of model membrane systems and structure determination methods. The code design ensures the geometries, including the pore, are compatible with different energy functions, as well as different applications, such as refinement, docking, and design. These applications can be utilized with experimentally determined structures as well as structures generated by other prediction algorithms such as AlphaFold2. In many cases this framework improves sampling and scoring of MPs.

## Methods

### Implementation of implicit membrane geometries

The RosettaMP framework has score functions that include physics-based score terms that depend on an atom’s position relative to the membrane. This dependence on the membrane is achieved through a function that models the transition from the hydrophobic, membrane phase to the aqueous phase, referred to as the transition function. The transition function is adapted from IMM1 [40]. The transition function ranges from zero to one, where an atom in the hydrophobic phase has a value of zero, and an atom far from the membrane has a value of one is scored as it is in an aqueous environment (Figure 1A).

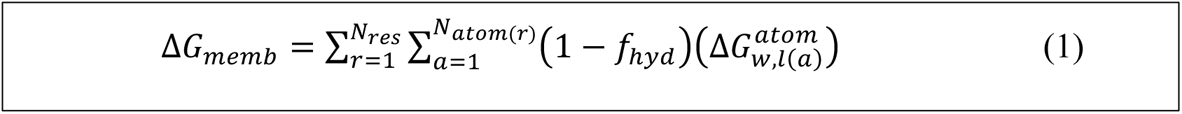

For example, in the `fa_water_to_bilayer` score term from *franklin2019* (equation 1) the 𝑓*_hyd_* is the transition function [39]. The 𝑓*_hyd_* transition function is a composition of two functions (equation 2),

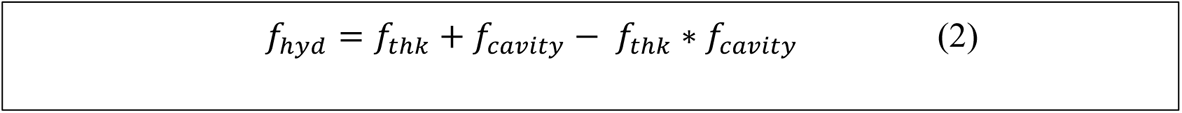

one representing the transition along the membrane normal and the other represents an aqueous pore as described in [48] and [39] (Figure 1B). In equation 2, 𝑓*_cavity_* describes the transition in the pore and 𝑓*_thk_* describes the transition out of the bilayer. Two other membrane dependent score terms, `fa_mpenv`

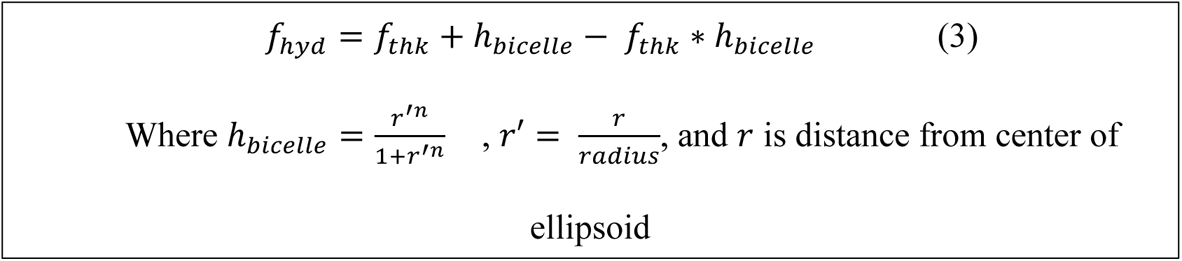

and `fa_mpsolv`, are based on the transition function used in IMM1 [31]. Therefore, the first step in implementing new geometries was to adapt the existing transition functions to accommodate different geometries. The transition function can be adapted to an ellipsoid geometry using a composition of functions (equation 3), similar to how the pore is modeled [48] (Figure 1C, D). In equation 3, the radius is what we refer to as the outer radius of the bicelle (Figure 1D). The user can set the inner radius using the -mp:geo:bicelle_radius option. The inner radius corresponds to the radius of the planar section of the ellipsoid, before the curve of the edges (Figure 1D). For a micelle with no planar section, the inner_radius is 0. The outer radius is set as the inner radius + the thickness of the membrane. The radius of micelles and bicelles in experiments depend both on the lipid and detergents used and the protein itself. Commonly used micelle and bicelle sizes have been studied and values or equations that depend on the concentration of detergent and lipid to calculate the radius have been reported [62, 63]. If the user does not set a bicelle radius, then one is estimated by calculating the largest distance found between two alpha carbons within three angstroms of the center of the membrane, dividing that distance in half and multiplying by three to set the bicelle inner radius. This ensures that if a user does not set a radius for the bicelle, that their protein is not modeled in a bicelle that is smaller than the protein itself which would not make physical sense, since the lipids or detergent molecules surround the protein so the size is dependent on the transmembrane domain of the protein [64]. If the user does provide a radius smaller than the largest distance between alpha carbons in the center of the membrane, a warning is printed that a larger radius may be needed. Adaptation of the transition function to a sphere is described by Nepal, Leveritt III, and Lazaridis [65] (Figure 1E). The origin continues to be at the center of the protein and in the middle of the membrane layers. The center of the sphere is at (0, 0, -radius) or (0, 0, +radius) depending on if the protein should be modeled with the membrane curved down (-radius) or the membrane curved up (+radius). The double bilayer geometry is simply a composition of two sphere equations (Figure 1F). The origin is at the center of the inner sphere. We have the user set the radius of the inner sphere and the distance between the inner and outer sphere, specifically the distance from the outer edge of the inner membrane to the inner edge of the outer membrane. To help the user know how their MP of interest is being scored in the implicit membrane environment we created an application that maps the transition function value for each atom into the b factor column of a PDB file.

To achieve our goals of flexible use across many applications and ease of further development, we implemented the new geometries as an interface class MembraneGeometry and each individual shape as an inherited class from MembraneGeometry (Figure S1). Within the RosettaMP framework the MembraneInfo class stores all necessary information for the implicit membrane [41]. Therefore, MembraneGeometry is a member of MembraneInfo, so anytime there is an instance of MembraneInfo it will contain an instance of MembraneGeometry. Score terms, such as FaMPEnv, FaMPSolv, and FaWaterToBilayerEnergy, that depend on an atom’s relative position to the membrane, the transition function value, get that information from MembraneGeometry. Previously, even for score terms that used the same transition function, that transition function was defined in the calculation for each score term. This creates a situation where one could be updated without updating all of them causing a mismatch in the implicit membrane being represented across score terms. This update ensures that the implicit membrane modeled is consistent across score terms.

The MembraneGeometry interface class houses members that are used across multiple geometries and virtual functions that must be defined in each inherited class. Thickness, steepness, and pore_parameters are variables that are needed across all geometries. Some geometry classes, such as Bicelle, have additional variables that are stored in the class itself, shown in Figure S1 in the top box of each geometry. There are three pure virtual functions contained in MembraneGeometry that must be defined in all inherited classes. These are shown in italics in the lower boxes for each class. The first pure virtual function is f_transition which returns the value of the transition function for an atom. The other virtual functions are 𝑓1 and 𝑓2. These are functions that allow for the calculation of the transition function derivative for gradient based minimization and are described in Abe et al [66] and this video: https://youtu.be/j07ibj-fT1A. For 𝑓1 and 𝑓2 one must first determine the point in space that the transition function value depends on the distance between the atom and that point, referred to as 𝑟*_alpha_*. For

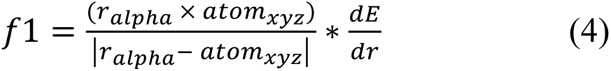

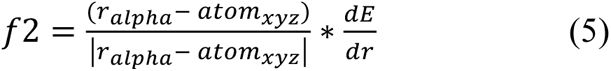

example, if the membrane plane is at z=0 and the membrane normal is parallel to the z-axis, for the slab model 𝑟*_alpha_* for an atom at (x1, y1, z1) 𝑟*_alpha_* is (x1, y1, 0). If the derivative of the transition function with respect to an atom’s distance from 𝑟*_alpha_* is 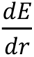, then 𝑓1 and 𝑓2 can be described as in equation 4 and 5. If the transition function depends on more than one distance, then 𝑓1 and 𝑓2 may need to contain multiple partial derivatives.

There are also methods used across multiple geometries that are defined in MembraneGeometry. These include the originally implemented transition function named f_imm1, the transition function for franklin2019 energy function named f_franklin, f_cavity, and f_hyd which returns the composition function to include the pore if there is a pore (Figure S1). Each child class also has methods that are specific for describing that geometry.

To ensure proper implementation of each derivative, we implemented unit tests utilizing the Rosetta benchmarking framework [67] checking that the analytical and numerical derivatives are in fact equal when calculated during minimization. After these tests were set up for each geometry, including the classic slab, we realized that some of the previous implementations of the derivatives needed to be updated. FaMPSolv energy term is a two-body term where the value depends both on an atom’s relative distance to the membrane center and at atom’s distance to another atom [31, 41]. However, the previous implementation only had the partial derivative with respect to the atom’s relative distance to the membrane center. We looked at how this derivative might impact results by running refinement on KCNE3 with the *mpframework2012* energy function that includes the FaMPSolv energy term (Figure S4). We see only minor shifts in score and RMSD distributions with the slab geometry before and after the derivative correction. Refinement in the bicelle geometry impacts the distribution the most, similar to the impact using the franklin2019 energy function (Figure 5I and L). The FaWaterToBilayer energy term did not include the partial derivative to the membrane center, but since the f_cavity depends on this distance it needed to be included as well.

### Protein Design with pore

The protein design analysis done is based on the MP Sequence Recovery benchmark test implemented within the Rosetta test server framework [51, 67]. Since we were specifically looking at how inclusion of the pore affected the sequence recovery, we used a subset of the original dataset of only MPs that included a pore [68]. We ran Rosetta’s fixed-backbone design [69] for each protein with both *franklin2019* and *mpframework2012* energy functions with and without the pore included. The pore functionality was turned off using the `-has_pore 0` flag. To note, by including this flag with either 0, 1, or empty, the pore is not calculated. We ran the fixed-backbone design protocol three times for each case. Residues were classified as pore-facing if 𝑓*_cavity_* > 0.1 and 𝑓*_thk_* < 0.75, from equation 2.

### Relax on Piezo 1

We used PDB ID 6B3R for the starting structure of Piezo 1, downloaded from the OPM database. The preparation and orientation of the structure is described in detail in the supplemental protocol capture. After orientation a spanfile was generated using the `mp_span_from_pdb` Rosetta application. We used the `mp_transition_bfactor` application to visualize the implicit membrane on the structure. We ran the FastRelax protocol using the franklin2019 energy function with either the slab or vesicle geometry with a radius of 120 angstroms.

### AlphaFold2 and Rosetta Relax on Mitochondrial TOM complex

We used AlphaFold2 multimer [53] to generate initial decoys for the mitochondrial TOM complex. The chain sequences from PDB ID 6UCU were used as input [52], with five multimer predictions per model, making a total of twenty-five models. A span file was created by downloading 6ucu from PPM 2.0 and running the `mp_span_from_pdb` Rosetta application. Output models from AlphaFold2 were transformed into membrane coordinates using `mp_transform` application, followed by deleting the EMB virtual residues in the output. These transformed structures were used as input to a FastRelax protocol using the franklin2019 energy function with either the slab geometry or a vesicle geometry with a radius of 180 Å, based on reported radius of curvature from PPM 3.0 [44]. For each of the twenty-five AlphaFold2 predictions, eighty structures were generated by the FastRelax protocol for a total of 2000 structures for each geometry. Note, to calculate the RMSD within Rosetta we had to edit the chain IDs to match the output from AlphaFold2.

### Relax on gap junction channel and AcrABZ-TolC multidrug efflux pump in double membrane geometry

The structure for the gap junction channel was obtained by downloading PDB ID 6MHQ [25] from the OPM webserver [45, 46]. A spanfile was created using the `mp_span_from_pdb` Rosetta application. The spanfile corresponds to the residues that span the inner membrane. The outer membrane is determined by setting the option `-mp:geo:double_vesicle_distance` to the distance from the outer edge of the inner membrane to the inner edge of the outer membrane. For this example, the double_vesicle_distance was set at 40 and the inner vesicle radius was set to 1000 Å, making the membrane practically flat. We ran FastRelax protocol using *franklin2019* to produce a total of 450 structures in both the slab and double vesicle geometries. The RMSD was calculated with respect to the original pdb structure 6MHQ.

The structure for the AcrABZ-TolC multidrug efflux pump was obtained by downloading PDB ID 5O66 [26] from the OPM webserver [46]. A spanfile was similar to above. The double_vesicle distance was set to 244 Å and the inner vesicle radius was 1000 Å. We ran FastRelax protocol using *franklin2019* to produce a total of 200 structures in both the slab and double vesicle geometries.

### Docking glycophorin A in micelle geometry

To dock glycophorin A we followed the steps described in Alford, Samanta, and Gray [51] for protein-protein docking using the *franklin2019* energy function. The structure for glycophorin A was downloaded from both the Protein Data Bank (PDB) [70, 71] and the OPM webserver [46]. The OPM website only provides the first conformation of the NMR ensemble, so we first wanted to look at all the conformations. However, we aligned to the OPM structure to move the structures into membrane coordinates as described in Alford et al. [41, 72]. We chose to use the first conformation in the NMR ensemble. We did both the prepacking and docking steps in the slab and bicelles geometries, so the geometry stayed consistent throughout the protocol. We created fifty models during the prepacking step and used the lowest scoring model as input into the docking step. We output 5000 models for the docking step. The bicelle inner radius was set at 1 Å.

### Relax KCNE3 in bicelle geometry

The NMR conformational ensemble PDB ID 2NDJ [57] was downloaded from the PDB and from the OPM webserver. A spanfile was created using the `mp_span_from_pdb` Rosetta application with the structure downloaded from the OPM web server. All the conformations from 2NDJ were transformed with this spanfile using the `mp_transform` Rosetta application. We chose to use the fourth conformation in the NMR ensemble since the amphipathic helix was position beside the transmembrane helix, where in the slab geometry it would be scored as if it was dipping into the membrane (Figure 5G). For both the slab and bicelles geometries 1000 output models were produced with the FastRelax protocol using *franklin2019*. The bicelles inner radius was set to 2 Å. The RMSD was calculated with respect to the starting structure, the 4^th^ conformation from PDB ID 2NDJ.

## Notes

### Competing Interest Statement

The authors have declared no competing interest.

